# ZEB2 transduces HIF1α dependent regulation of Transglutaminase 2 in glomerular podocytes

**DOI:** 10.1101/2021.12.15.472753

**Authors:** Lakshmi P Kolligundla, Ashish K Singh, Rajesh Kavvuri, Anil K Pasupulati

## Abstract

Glomerular podocytes are instrumental in ensuring glomerular permselectivity and regulating the integrity of glomerular biology. However, podocytes are vulnerable to various noxious stimuli such as hypoxia, and podocyte injury presented with glomerulosclerosis and impaired kidney function. The mechanism of hypoxia-induced podocyte injury vis-a-vis glomerulosclerosis has remained enigmatic. Hypoxia inducible factor 1α (HIF1α) that transduces hypoxic adaptations, induces Transglutaminase 2 (TG2), a calcium dependent enzyme that catalyzes intramolecular ε-(γ-glutamyl) lysine cross-links of extracellular matrix (ECM) proteins. In this study, we investigated the mechanism of regulation of TG2 by HIF1α. Stabilization of HIF1α by FG4592 (Roxadustat) and physiological hypoxia, resulted in elevated expression of ZEB2 (zinc-finger E-box-homeobox 2) and its downstream target TRPC6 (transient receptor potential channel 6). ZEB2 transcriptionally activates TG2 expression, whereas, via TRPC6, it induces calcium influx, inturn it increases the TG2 activity. Blocking the TRPC6 action or suppressing its expression only partially attenuated FG4592 induced TG2 activity, whereas suppression of ZEB2 expression significantly abolished TG2 activity. This study demonstrates that stabilization of HIF1α stimulates both TG2 expression and activity, whereas abrogation of HIF1α by metformin prevented HIF1α regulated TG2 and consequent glomerular injury.

## Introduction

Glomerular diseases account for most of the chronic kidney diseases (CKD), which eventually progresses to end-stage kidney failure (ESKF) (1). CKD affects about one in every ten people in the USA, and patients with CKD have a 3 to 5 fold increased risk of mortality (2) . The progressive nature of CKD is associated with a persistent loss of tissue and its replacement by extracellular matrix (ECM), culminating in kidney fibrosis. Although ECM provides structural and biological support to neighboring cells, uncontrolled accumulation and excess deposition of ECM protein leads to the progressive loss of kidney architecture and compliance.

The function of the vertebrate kidney to maintain homeostasis is attributed to its smallest functional unit, the nephron, which comprises two parts; glomerulus and tubule. While the glomerulus ensures the ultrafiltration of blood and the formation of primary urine, the tubular part involves reabsorption and secretion; thus, both glomerulus and tubule work in concert and determine the final urine composition. The glomerular perm-selectivity is contributed by a three-layered glomerular filtration barrier (GFB) that allows the filtration of molecules are smaller than 60 kDa (3). GFB comprises of inner fenestrated capillary endothelial cells, middle glomerular basement membrane (GBM), and outer podocyte foot-processes (4). The podocyte is a terminally differentiated glomerular visceral epithelial cell with primary and secondary foot processes that enwrap the capillaries and offer epithelial coverage (5). Podocytes account for about 30% of glomerular cells and secrete vascular endothelial growth factor (VEGF) and ECM components to help maintains the endothelium and GBM, respectively. Since podocytes are instrumental for glomerular filtration, podocyte injury induces a complex set of biological responses in the glomeruli, leading to proteinuria (6). Glomerular diseases, such as minimal change disease, diabetic nephropathy, and focal segmental glomerulosclerosis (FSGS) begin with podocyte injury and often end up with fibrotic phenotype, which is initiated by mesangial cell activation, ECM overproduction, and scar formation (1,6).

The homeostasis of ECM is regulated by specialized enzymes, transglutaminases (7). The normal function of transglutaminases is to cross-link proteins that are involved in several biological processes ranging from tissue repair to strengthening the fertilization envelope (8). Transglutaminase 2 (TG2; EC: 2.3.2.13), a member of the transglutaminase family, is majorly associated with ECM cross-linking with the capacity to catalyze acyl transfer reaction between protein-bound glutamine residue and protein-bound lysine residue (cross-linkage) or poly-amination (9). The TG2 enabled ε(γ-glutamyl) lysine iso-peptide cross-linking facilitates inappropriate deposition of ECM proteins and is resistant to proteolytic degradation (10). Several ECM proteins (eg: Collagen) are substrates of TG2 (11). Although TG2 is implicated in scarring of liver (12) and lung (13) the enzyme has been extensively characterized in kidney fibrosis (14). A strong correlation between levels of TG2 (R2 = 0.92), ε(γ-glutamyl) lysine iso-peptide cross-linking (R2 = 0.86), and the development of tissue scarring was reported (14). Since TG2 is considered a therapeutic target to combat fibrotic diseases, including kidney fibrosis and CKD, understanding the mechanism of TG2 regulation is of great importance. Kidney fibrosis is considered as terminal pathway involved in the continuous progression of CKD which is characterized by glomerulosclerosis and tubulointerstitial fibrosis (15). Since hypoxia is considered to be one of the most common causes of kidney fibrosis and TG2 transduces fibrotic events, the regulation of TG2 expression and its activity in hypoxic condition needs to be elucidated. As hypoxia inducible factor 1α (HIF1α) elicits cellular effects of hypoxia, we investigated the mechanism of regulation of TG2 in the settings of elevated HIF1α and found that HIF1α /ZEB2 axis controls both TG2 expression and activity.

## Results

### Co-expression of HIF1α and TG2 in glomerular podocytes

Data mining using Nephroseq (https://www.nephroseq.org), (University of Michigan, Ann Arbor, MI) revealed co-expression of HIF1α, ZEB2 (Zinc finger E-box-binding homeobox 2), TRPC6 (Transient receptor potential channel 6), and TG2 in Nakagawa CKD validated dataset (Fig.1A). To assess the specific role of HIF1α on TG2 expression, we treated podocytes with FG4592 (known as Roxadustat, a prolyl hydroxylase inhibitor) in both dose (1-15uM; Fig.1B) and time dependent manner (3-24h; Fig.1C). Stabilization of HIF1α using a prolyl hydroxylase inhibitor resulted in elevated expression of TG2 (Fig.1B&C). Expression of TG2 in FG4592 treated podocytes was demonstrated by immunofluorescence (Fig.1D). Furthermore, we validated the Nephroseq data and found that ZEB2, TRPC6, and TG2 expression was elevated in podocytes treated with FG4592 (Fig.1E&F).

**Figure 1.**
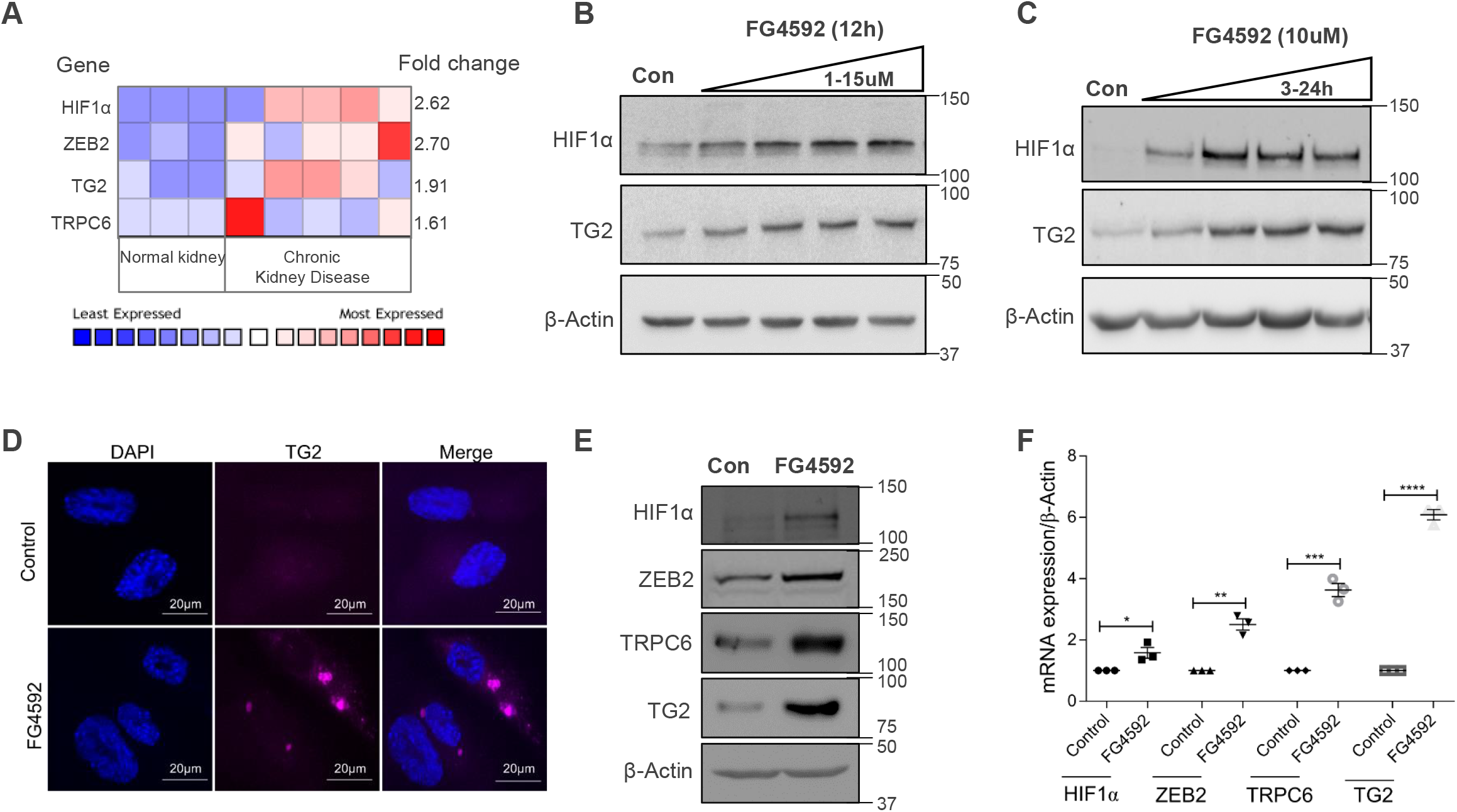
HIF1α accumulation is associated with TG2 expression in podocytes. **(A)** Expression of *HIF1α, ZEB2, TG2*, and *TRPC6* was analyzed between healthy volunteers (n=3) and chronic kidney disease patients (n=5) in the validated set group of Nakagawa CKD kidney database in Nephroseq (www.nephroseq.org). Immunoblotting analysis of HIF1α, TG2, and β-actin in human glomerular podocytes treated with **(B)** 1,5,10, and 15 µM of FG4592 for 12h and (**C**) 10µM FG4592 for 3,6,12, and 24h. **(D)** Immunofluorescence detection of TG2 in control vs. FG4592 treated podocytes. Scale bar 20μm. Nephroseq data was validated in human podocytes exposed to 10µM of FG4592 for 24h by **(E)** immunoblotting and (**F**) qRT-PCR. Error bars indicate mean ± SE; n=3. *, *p* < 0.02; **, *p* <0.001; ***, *p* < 0.0002; ****, *p* < 0.0001 by student t-test.

### Elevated intracellular calcium levels associate with increased TG2 activity

TG2 is a calcium-dependent enzyme. Since we observed co-expression of TRPC6, a nonselective cation channel and a component of store-operated calcium entry, we assessed calcium levels and TG2 activity in FG4592 treated podocytes. We measured intracellular calcium using calcium-sensitive Fluo-3 AM dye and observed elevated calcium levels in podocytes treated with FG4592 (Fig.2A). Podocytes treated with a membrane-permeable 2-aminoethoxydiphenyl borate (2-APB), an inhibitor of TRP channels abrogated FG4592 induced calcium flux into podocytes (Fig.2A). Alternatively, FG4592 induced calcium influx was attenuated in podocytes in which TRPC6 expression was knocked down (Fig.2A). We next assessed TG2 activity in podocytes in which TRPC6 expression was attenuated by transfecting siRNA, or TRPC6 activity was inhibited with 2-APB. Increased TG2 activity was observed with FG4592 treatment, only partially reduced when podocytes either co-treated with 2-APB or siTRPC6 (Fig.2B). We next determined TRPC6 and TG2 levels in podocytes transfected with siTRPC6 by immunoblotting. Control was transfected with scrambled RNA (Fig.2C&D). The data suggest reduced expression of TRPC6 by siRNA did not wholly abolish TG2 expression in FG4592 treated cells; this could explain the residual TG2 activity in podocytes in which TRPC6 expression was knocked down. These observations suggest that, though increased intracellular calcium levels via TRPC6 elicits TG2 activity, elevated TG2 expression during hypoxia settings could further contribute to the increased TG2 activity.

**Figure 2.**
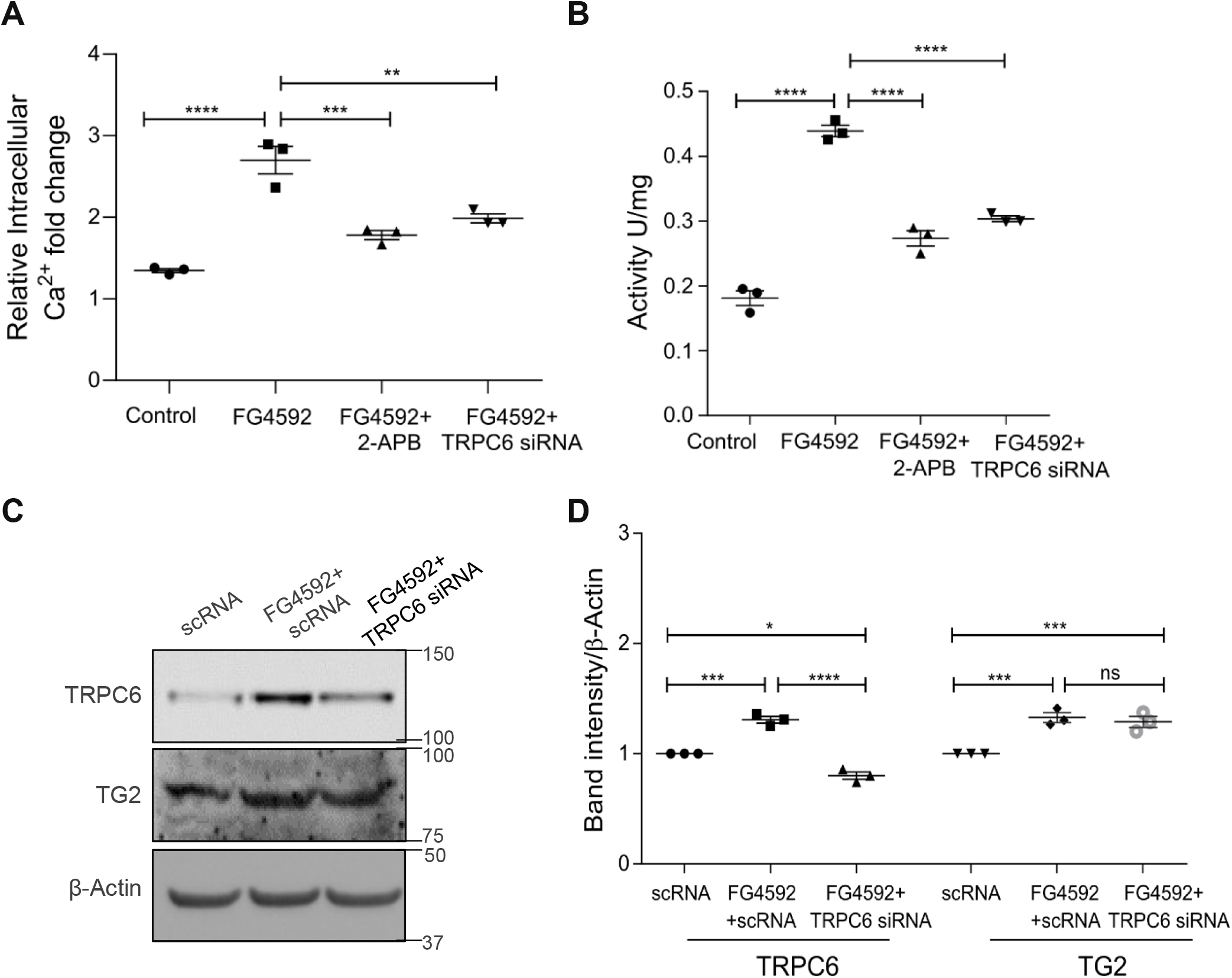
HIF1α elevates TG2 activity by inducing the calcium influx. **(A)** Enhanced calcium influx was observed in FG4592 treated podocytes whereas 2-APB, a calcium channel blocker; and TRPC6 knockdown abrogated FG4592 induced calcium flux. Error bars indicate mean ± SE; n=3. **, *p* < 0.001; ***, *p* < 0.0005; ****, *p* < 0.0001 by one-way ANOVA after Tukey’s multiple comparison test. (**B**) TG2 activity was measured as a readout of hydroxamate formed in the invitro TG2 activvity in podocyte lysates from FG4592, 2-APB, and TRPC6 knockdown conditions and the activity represented as units per mg of protein. Error bars indicate mean ± SE; n=3. ****, *p* < 0.0001 by one-way ANOVA after Tukey’s multiple comparison test. **(C)** TRPC6 and TG2 expression was assessed in podocytes transfected with TRPC6 siRNA and treated with or without FG4592. **(D)** Densitometric analysis of western blots was normalized to β-actin. Data presented as mean ± SE; n=3. *, *p* < 0.01; ***, *p* < 0.0005; ****, *p* < 0.0001 by one-way ANOVA after Tukey’s multiple comparison test.

### Essential role of ZEB2 in HIF1α induced TG2 expression and activity

Since we observed TRPC6 knockdown does not completely eliminate TG2 activity despite reduced intracellular calcium levels, we hypothesize that TG2 expression is regulated differentially during hypoxia. It was shown earlier that ZEB2 is a bona fide target of HIF1α, and ZEB2 expression was induced in ischemic conditions (16). We investigated the mechanistic insights since we observed co-expression of ZEB2 and TG2 in the Nephroseq database and podocytes exposed to FG4592 (Fig.1A&E&F) of this temporal association. We found three ZEB2 binding sites (E-box-binding region) upstream to the transcription start site (TSS) of TG2 in the range of - 202 to -207; -238 to -243; -309 to -314 proximal promoter region of TG2 (Fig.3A). The promoter sequence of TG2 got from the Eukaryotic Promoter Database (EPD) and ZEB2 Transcription factor sequence was obtained from Okamoto et al (17). Further, we performed a ChIP assay that revealed the binding of ZEB2 to the TG2 promoter region (Fig.3B). N-cadherin promoter was used as a positive control as it possesses a putative ZEB2 binding site, and IgG acts as a negative control. Ectopic expression of ZEB2 resulted in elevated levels of both TRPC6 and TG2 in HEK293T cells, and in contrast, knockdown of ZEB2 abolished TRPC6 and TG2 expression (Fig.3C). Interestingly, ectopic expression of ZEB2 resulted in elevated intracellular calcium levels, while ZEB2 knockdown manifested in decreased intracellular calcium levels (Fig.3D), which is in tandem with TRPC6 expression. We next assessed TG2 activity in podocytes that ectopically express ZEB2 and in podocytes where ZEB2 expression was knocked down (Fig.3E). Increased TG2 activity was observed with FG4592 treatment as well as in ZEB2 over expressed (OE) condition (Fig.3E). On the other hand, attenuation of ZEB2 expression resulted in diminished TG2 activity (Fig.3E). Together, the data suggest ZEB2 regulates TG2 expression directly and its activity via inducing TRPC6 dependent calcium flux.

**Figure 3.**
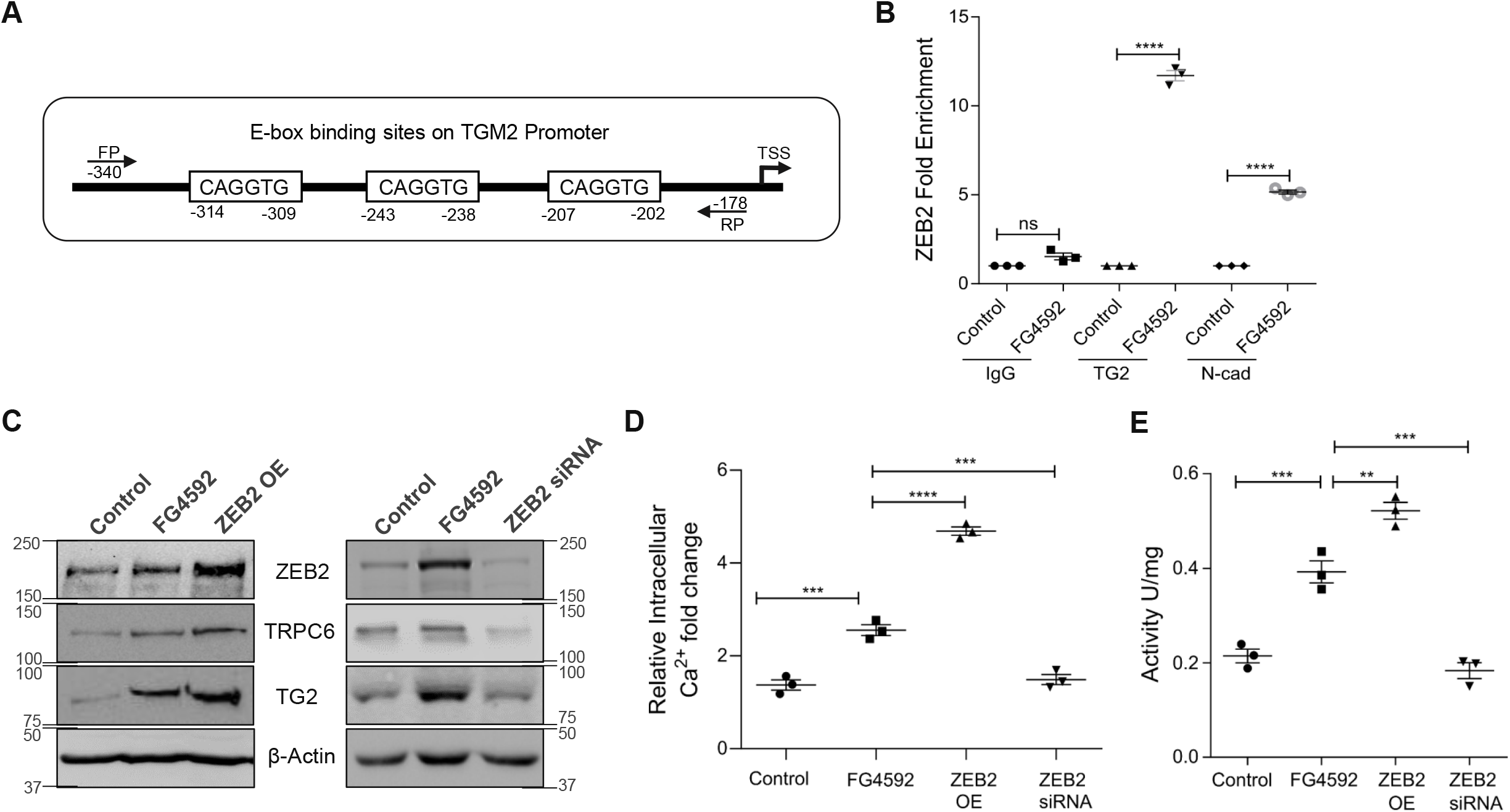
ZEB2 regulates the TG2 expression directly and activity via TRPC6. **(A)** The promoter region of TG2 with ZEB2 binding E-box regions and region that was amplified following ChIP is indicated by arrows. **(B)** DNA from immunoprecipitated samples of podocytes treated with or without FG4592 was PCR amplified for ZEB2. E2-box in N-cadherin promoter serves as a positive control for ZEB2 interaction. Data presented as mean ± SE; n=3. ****, *p*< 0.0001 by one-way ANOVA after Tukey’s multiple comparison test. **(C)** Immunoblotting analysis showing protein levels of ZEB2, TRPC6, and TG2 in HEK293T cells which are ectopically express ZEB2 and ZEB2 knockdown conditions. **(D)** Intracellular calcium flux was observed in FG4592 treated podocytes and podocytes in which ZEB2 is ectopically expressed or knockdown. Data presented as mean ± SE; n=3. ***, *p* < 0.0005; ****, *p* < 0.0001 by one-way ANOVA after Tukey’s multiple comparison test. **(E)** TG2 activity was assessed in podocyte lysate treated with FG4592 and podocytes in which ZEB2 is ectopically expressed or knockdown. Data presented as ± SE; n=3. **, *p* < 0.001; ***, *p* < 0.0005; by one-way ANOVA after Tukey’s multiple comparison test.

### Metformin suppresses HIF1α dependent TG2 expression both in vitro and in vivo

Since we observed stabilization of HIF1α resulted in both enhanced TG2 expression and activity, we assessed whether preventing HIF1α accumulation does affect TG2 expression and activity. In a recent study, Hart et al demonstrated that metformin activates prolyl hydroxylases and ensure the degradation of HIF1α in mesothelial cells (18). Metformin was shown to reverse hypoxia-induced migration by targeting the HIF1α/VEGF pathway in gall bladder cancer cells (19). TG2 has been postulated as an inducer for epithelial–mesenchymal transition (EMT), which is responsible for the fibrosis advancement in CKD (20,21). FG4592 induced stabilization of HIF1α and consequent accumulation of TG2 and mesenchymal markers such as vimentin, fibronectin, and α-smooth muscle actin (α-SMA) were attenuated in podocytes that were treated with metformin (Fig.4A). Metformin prevented HIF1α accumulation in podocytes that were exposed to physiological hypoxia (1% oxygen) (Fig 4B). Metformin also prevented accumulation of down-stream targets of HIF1α such as TG2 and mesenchymal markers in podocytes exposed to hypoxia (Fig. 4B). Similarly, HIF1α inhibitor abolished the hypoxia induced stabilization of HIF1α, TG2 and mesenchymal markers (Fig. 4C). Immunofluorescence data also revealed the reduced HIF1α and TG2 expression in podocytes that were treated with metformin (Fig.4D&4E). Together, the data suggests that metformin prevents HIF1α accumulation and abolishes hypoxia induced activation of ZEB2/TRPC6/TG2 axis in podocytes.

**Figure 4.**
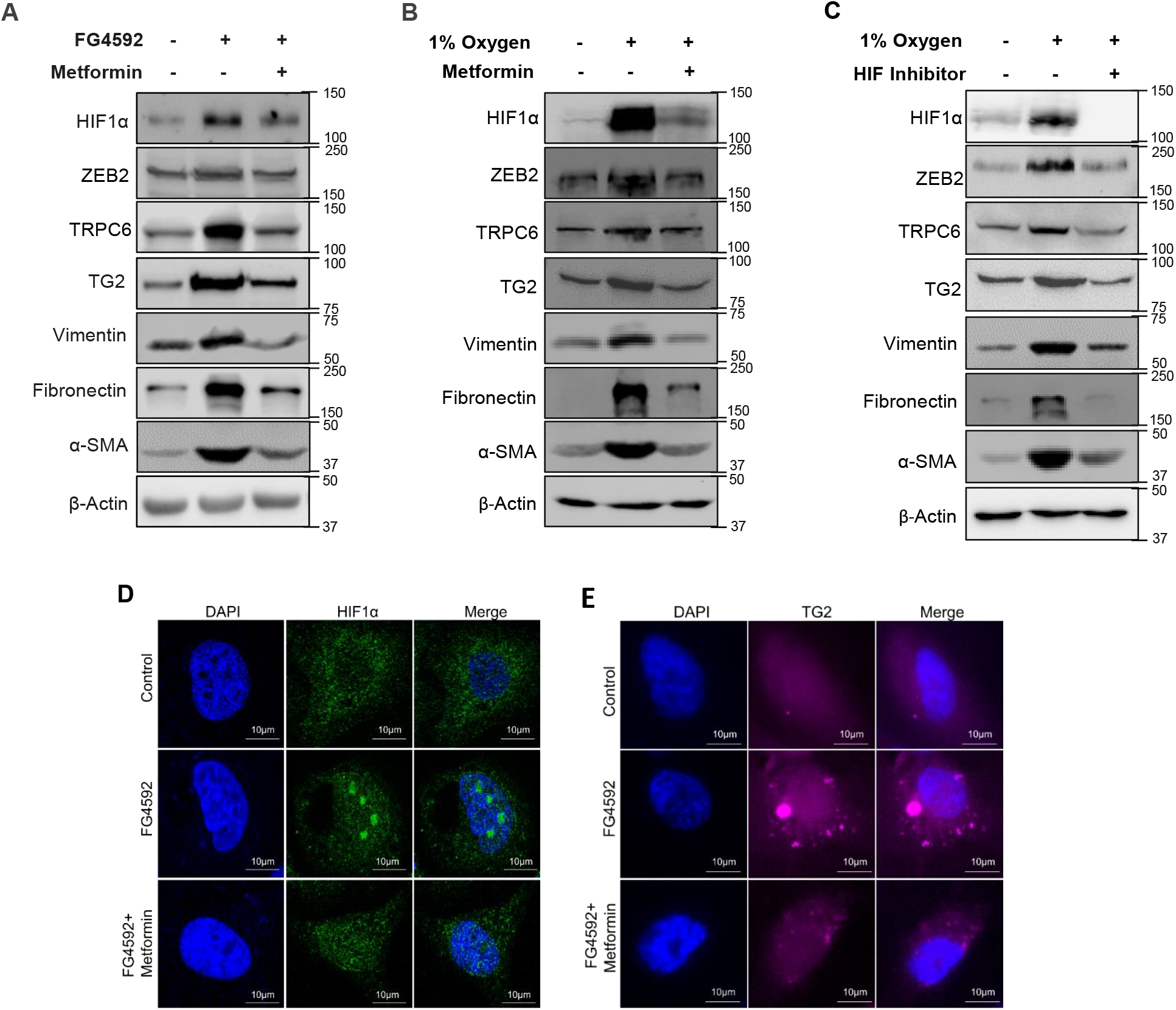
Metformin abrogates ZEB2 induced TG2 expression. Immunoblotting analysis of HIF1α, ZEB2,TRPC6,TG2, vimentin, fibronectin, and α-SMA in podocytes **(A)** treated with FG4592 in the presence or absence of metformin or **(B)** exposed to hypoxia (1% oxygen) in the presence or absence of metformin, and **(C)** exposed to hypoxia (1% oxygen) in the presence or absence of HIF1α inhibitor. Immunofluorescence detection of **(D)** HIF1α and **(E)** TG2 in podocytes treated with FG4592 with or without metformin treatment. Scale bar 10μm.

### Metformin ameliorates renal fibrosis and prevents proteinuria

Further, we investigated the effect of stabilization of HIF1α on TG2 in vivo. Similar to in vitro studies, we observed elevated expression of HIF1α, ZEB2, TRPC6, TG2, and mesenchymal markers in glomerular lysate from mice administered with FG4592 (5mg/Kg b.w for 3 months) whereas co-administration of metformin (250mg/Kg b.w) prevented activation of ZEB2/TRPC6/TG2 axis and attenuated expression of mesenchymal markers (Fig.5A). We assessed the histological features by Masson’s trichrome, PAS, and H&E staining in kidney sections from mice treated with FG4592 with or without metformin (Fig.5B). Glomerular injury score analysis revealed metformin attenuated FG4592 induced fibrosis and altered histological features in glomerular and periglomerular regions as evidenced by glomerular injury score (Fig.5C). Fibrosis causes the stiffening of basement membrane that counteracts podocyte interactions with GBM leading to retraction of foot processes, called foot process effacement (FPE). To study the FPE, we performed transmission electron microscopy analysis in mice injected with FG4592 with or without metformin. Our results show that metformin protected the mice from FG4592 induced podocyte FPE (Fig.5D).

**Figure 5.**
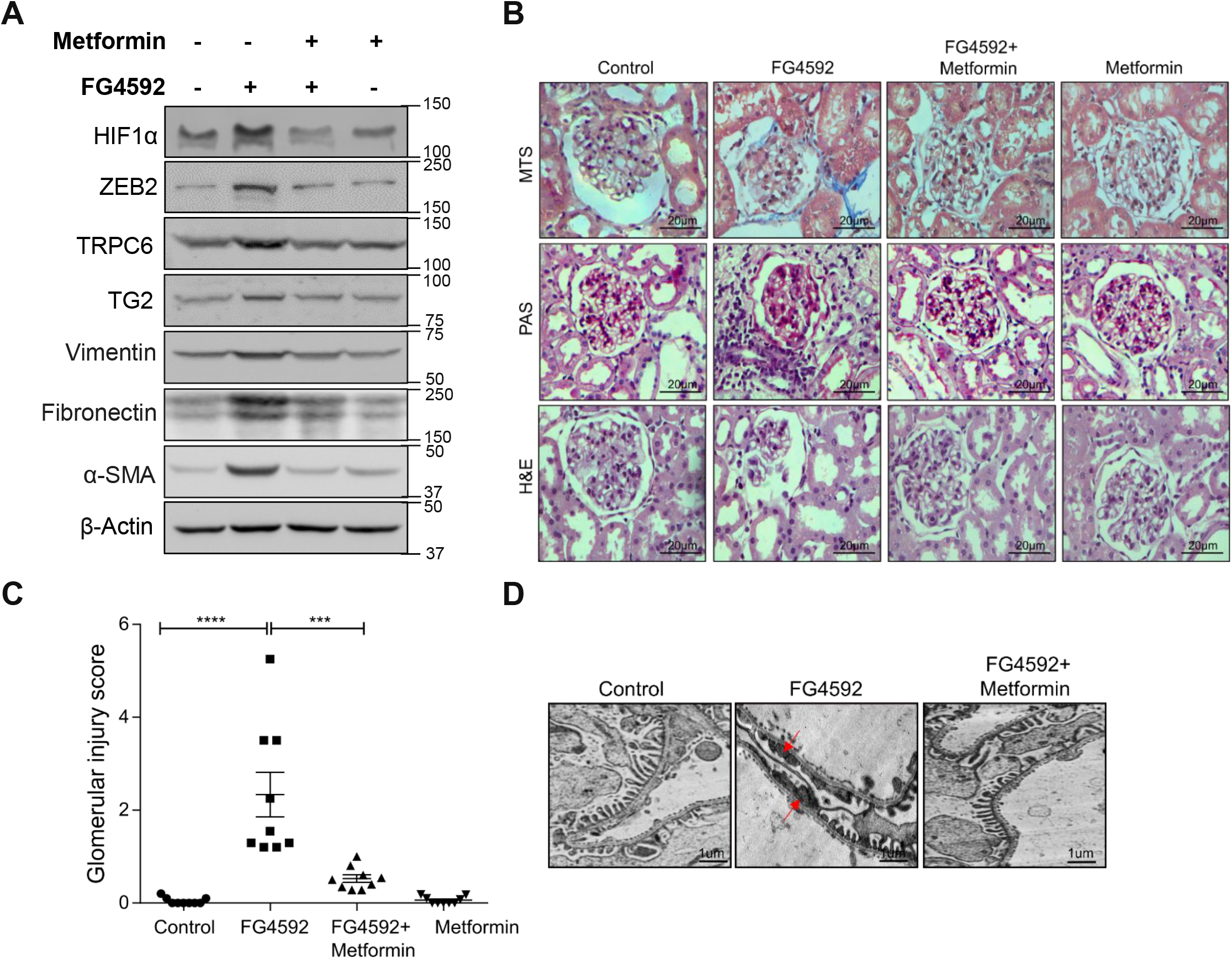
Metformin prevents hypoxia-induced glomerulosclerosis. **(A)** Immunoblotting analysis of HIF1α, ZEB2, TRPC6, TG2 and mesenchymal markers in glomerular lysate from mice administered with FG4592 and treated with or without metformin. **(B)** Masson’s Trichrome Staining (MTS), PAS, and H&E staining of glomerular regions of mice administered with FG4592 and treated with or without metformin. Scale bar 20μm. **(C)** Glomerular injury analysis was performed as described in the methods section. Data presented as mean ± SE; n=9. ***, *p* < 0.0005; ****, *p* < 0.0001 by one-way ANOVA after Tukey’s multiple comparison test. **(D)** TEM images of podocytes from mice administered with FG4592 and treated with or without metformin. Arrow marks indicate the podocyte FPE. Scale bar 1μm.

Metformin administration improved kidney function in FG4592 treated mice as evidenced by attenuation of proteinuria (Fig.6A) and urinary albumin to creatinine ratio (UACR) (Fig.6B). Since hypoxia is considered to be involved in renal pathology of patients with diabetes (22), we assessed the expression of ZEB2 and TG2 expression in patients with diabetes. Elevated expression of TG2 and ZEB2 expression was observed in glomerular sections from diabetic nephropathy patients (Fig. 6C). Albumin levels was estimated in the human diabetic nephropathy patient’s urine (Fig. 6D). Together these results suggest that attenuation of HIF1α prevent renal fibrosis and FPE in FG4592 treated mice, thus protect form proteinuria. Schematic illustration of metformin effect on FG4592 treated podocytes (Fig.6E).

**Figure 6.**
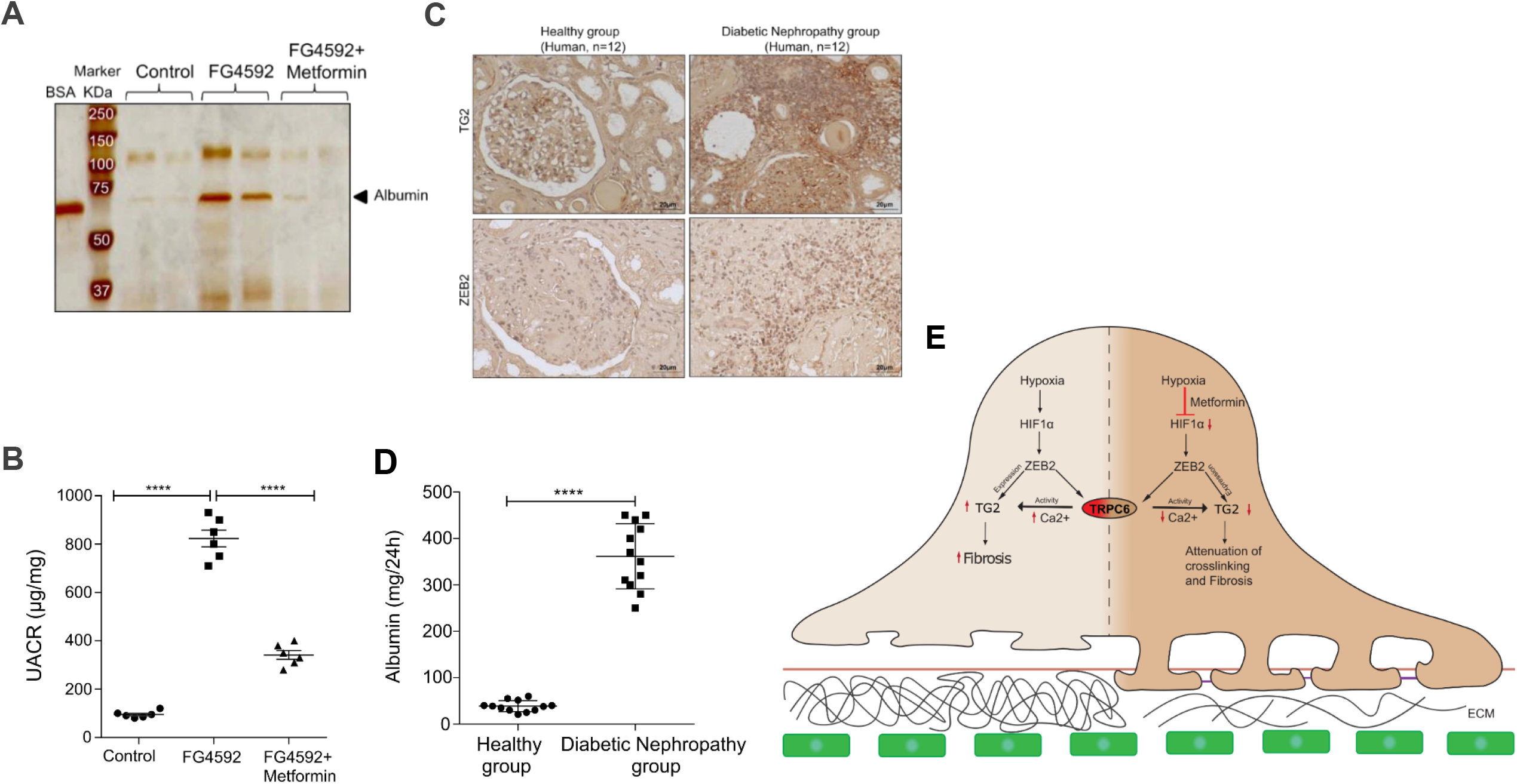
Metformin improves hypoxia-induced proteinuria. **(A)** Silver staining of urinary proteins from mice administered with FG4592 and metformin. **(B)** UACR from mice administered with FG4592 and treated with or without metformin. Data presented as mean ± SE; n=6. ****, *p* < 0.0001 by one-way ANOVA after Tukey’s multiple comparison test. **(C)** Representative images of immunohistochemical staining for TG2 and ZEB2 in glomerular sections from people with or without DN. Scale bars indicate 20μm. **(D)** Albumin levels were estimated from the human urine samples in healthy vs DN patients. Data presented as mean ± SE; n=12. ****, *p* < 0.0001 by student t-test. **(E)** A scheme depicting the mechanism of regulation of TG2 by HIF1α/ZEB2 axis.

## Discussion

Excessive deposition of ECM proteins and resistance to breakdown caused by cross-linking of ECM components is the predominant reason for fibrosis. It has been known that TG2 plays a vital role in cross-linking matrix proteins. In this study, we investigated the mechanism of TG2 regulation during hypoxia and in conditions that mimic hypoxia. Stabilization of HIF1α by FG4592 and 1% oxygen conditions resulted in elevated expression of ZEB2 and its downstream target TRPC6. ZEB2 transcriptionally activates TG2 expression, whereas, via TRPC6, it induces calcium influx, thus increase TG2 activity. Blocking the TRPC6 action or suppressing its expression only partially attenuated FG4592 induced TG2 activity, whereas suppression of ZEB2 expression significantly abolished TG2 activity. This study demonstrates that stabilization of HIF1α stimulates both TG2 expression and activity, whereas abrogation of HIF1α by metformin prevented HIF1α regulated TG2 and consequent glomerular injury (Fig. 6E).

HIF1α plays a crucial role in orchestrating the adaptive response of cells to the hypoxic condition by transcriptional activation of over a hundred downstream targets. The HIF1α targets regulate several vital biological processes such as erythropoiesis, angiogenesis, cell survival, cell metabolism, and EMT. Hypoxia/HIF1α signaling was shown to elicit several detrimental effects on podocyte morphology and function (23). ZEB2, an E-box binding homeobox 2 transcription factor, was reported to be a HIF target gene, and it induces TRPC6 expression (16). In this present study, we identified that ZEB2 regulates TG2 expression directly. TG2 promoter has E-box regions, and interaction of ZEB2 with TG2 promoter is evidenced by chromatin immunoprecipitation, whereas ZEB2 knockdown significantly abolished TG2 activity. On the other hand, inhibition of TRPC6 activity did not altogether abolish TG2 activity. This study reports the complex and multi-level regulation of TG2 by the ZEB2/TRPC6 axis in hypoxic conditions.

Besides catalyzing protein cross-linking and polyamination, TG2 is a critical mediator in regulating the gene expression at multiple levels. Increased TG2 expression is associated with the acquisition of EMT and stem cell-like characteristics in metastatic tumors found in lymph node and drug-resistant cancers (24,25). TG2 expression was associated with proteinuria and progression of immunoglobulin (Ig) A nephropathy (26). TG2 regulates gene transcription by modulating the activity of NF-κB (27,28). Earlier reports showed that TG2 regulates protein quantity by modifying and sequestrating proteins such as BAX-binding protein, caspase 3, and nucleophosmin (27,29,30). In a recent study it was showed that TG2 is involved in the translational control of mRNAs in response to hypoxic stress by increasing mTORC1-mediated phosphorylation of 4EBP1 and cap-dependent translation (31). Co-localization of TG2 and endostatin in the ECM secreted by endothelial cells under hypoxia shown to regulation of angiogenesis and tumorigenesis by the two proteins (32).

Podocytes are instrumental for the integrity of GFB. The proteinuria is characterized by the loss of podocytes, which leads to glomerulosclerosis and end-stage renal failure. The loss of more than 20% of podocytes causes irreversible glomerulosclerosis and a decrease in glomerular filtration rate (33,34). Yamaguchi et al proposed that EMT as a novel mechanism for podocyte loss (35). EMT is thought to play a role in tubulointerstitial fibrosis and renal disease progression (36,37). TG2 is a key component of the pro-inflammatory response, and it has been linked to EMT in both fibrosis and cancer. In a recent study, Shinde et al, reported that TG2 facilitates extracellular vesicle mediated epithelial-mesenchymal plasticity and metastatic niche of breast cancer cells (38). These evidences suggests that targeting TG2 could be a practical approach to combat EMT and metastasis or dealing with hypoxia-induced adverse manifestations. Therefore an adequate understanding of TG2 regulation helps in dealing with clinical conditions in which TG2 is implicated.

Earlier reports showed increased TG2 activity and expression in hypoxic conditions. TG2 expression was significantly up-regulated in chronic hypoxic rats and associated with right ventricular hypertrophy (39), whereas elevated TG2 activity is associated with hypoxia-induced pulmonary hypertension (40). In contrast, studies showed that TG2 is upstream of HIF1α and regulates the expression of HIF1α and its cellular targets. TG2 interacts with p65/RelA, and this complex binds to the HIF1α promoter and induces its transcriptional activation (41). Interestingly, inhibition of TG2 abolished HIF1α expression and attenuated ZEB2 expression (41). Though it is debatable whether TG2 is upstream or downstream of HIF1α, ZEB2 appears to be a transcriptional target of HIF1α (16,42,43).

ZEB2 is a SMAD-interacting transcription factor involved in Mowat-Wilson syndrome, a congenital disorder with an increased risk for kidney anomalies (44). Although studies reported that ZEB2 regulates the morphogenesis of mesenchyme-derived nephrons and is required for normal nephron development in mice (44), elevated expression of ZEB2 was implicated in the EMT of podocytes by suppressing E-cadherin expression (45). ZEB2 induction is associated with elevated expression of transforming growth factor-beta-induced (TGFBI) protein in growth hormone (GH) treated podocytes and proteinuria in rats administered with GH (46). TGFBI is an integral component of ECM, and it interacts with ECM proteins such as integrins and induces stabilization of microtubules (47). Cerebral ischemia induces the HIF1α/ZEB2 axis in the glomerular podocytes and contributes to proteinuria (16). The deleterious effects of the HIF1α/ZEB2 axis attributed to increased TRPC6 expression and resultant calcium influx into podocytes that resulted in aberrant activation of focal adhesion kinase (16). Therapeutically inhibiting ZEB2 expression or TRPC6 expression is not a suggested option. Inhibition of ZEB2 expression suppresses TRPC6 expression and calcium conductance into a cell. It was recently reported that decreased TRPC6 expression in TRPC6 –/– podocytes result in mislocalization of calpain and significant down-regulation of calpain activity, which resulted in altered podocyte cytoskeleton, motility, and adhesion (48). Therefore, blocking HIF1α accumulation or targeting TG2 active site inhibitors appears to be therapeutic options.

## Materials and Methods

### Materials

Anti-TG2 (NBP2-54633), anti-HIF1α (NB100-105), and anti-ZEB2 (NBP1-82991) antibodies were purchased from Novus Biologicals (Centennial, Colorado). Anti-fibronectin (#PAA037Hu01), anti-vimentin (#PAB040Hu01) were purchased from Cloud-clone (Houston, USA), anti-α-SMA (#19245), anti-actin (# 4970), and ChIP grade Protein-G agarose beads were purchased from Cell Signaling Technology (Danvers, MA). Anti-TRPC6 (#PA5-20256), RPMI 1640, DMEM, FBS, 100X antibiotics, and ProLong™ Diamond Antifade Mountant were purchased from Thermo Fisher Scientific (Waltham, MA). Secondary antibodies were purchased from Jackson laboratories (Bar Harbor, ME). TRIzol reagent were purchased from Life technologies (Carlsbad, CA) and cDNA reverse transcription kit obtained from Bio-Rad. Fluorescence based Alexa labeled secondary antibody (# A32728) were obtained from Invitrogen. Nitrocellulose membrane and ECL reagent, and SYBR Green Master Mix were obtained from Bio-Rad Laboratories (Hercules, CA). Metformin (# PHR1084), Fluo-3AM (#39294), 2-APB (#D9754), 5-BP (#914134), Z-gln-gly (#C6154), L -Glutamic acid γ-monohydroxamate (#G2253), X-tremeGENE 9 DNA Transfection Reagent and X-tremeGENE siRNA Transfection Reagents were purchased from Sigma-Aldrich (St. Louis, MO). FG4592 (#HY-13426) were purchased from MedChemExpress, India. TRPC6 siRNA (#SC-42672), ZEB2 siRNA (#SC-38641) and HIF1α inhibitor (#sc-205346) were purchased from Santa Cruz Biotechnology (Dallas, TX). Primers used in this study procured from Integrated DNA Technologies (Coralville, IA). All other reagents used were of analytical grade and obtained from Sigma Aldrich (St. Louis, MO, USA).

### Cell culture & *In vitro* experimental conditions

Human podocytes were obtained from Prof. Moin A Saleem (University of Bristol, Bristol, UK) and were cultured as described earlier (42,43). Podocytes were cultured at growth permissive conditions (33°C) and to induce differentiation these cells were shifted to 37°C and maintained for 14 days. Differentiated podocytes were used in all experimental procedures. 10mM stock of FG4592 was prepared in DMSO (treated with 10µM final concentration) and 1M stock of metformin was prepared in PBS (treated with 1mM final concentration). Experiments involved treatment of both compounds, metformin was first treated for 3h and then treated with FG4592. 30 mM stock of HIF1α inhibitor (methyl-3-[[2-[4-(2-adamantyl)phenoxy]acetyl]amino]-4-hydroxybenzoate) was prepared in DMSO (treated with 30µM final concentration). Experiments involved in FG4592 and HIF1α inhibitor both compound, HIF1α inhibitor was first treated with for 1h and then exposed to 1% oxygen. HEK293T cells were cultured in Dulbecco’s modified Eagle’s medium supplemented with 10% FBS with antibiotic & antimycotics and incubated at 37°C and 5% CO_2_.

### Immunoblotting

Equal amount of total podocyte cell lysate or glomerular lysate was subjected to SDS-PAGE and immunoblotting as described earlier (49). Band intensities were quantified using image J (NIH).

### Gene expression analysis

Total RNA was extracted from podocytes and gene expression analysis was performed as detailed earlier (50).

### Immunofluorescence

Podocytes were cultured on coverslips up to 60% confluence followed by treatment with FG4592 and metformin. After treatment those were fixed with 4% paraformaldehyde, permeabilized with 0.1% Triton X-100 and blocked with 3% bovine serum albumin (BSA) in 1X PBS. Cells were incubated with TG2 antibody (1:100 dilution) and followed by Alexa labeled secondary antibody (1:200 dilution). Mounting was done by using the Prolong gold antifade DAPI. Images were acquired using a trinocular microscope (Leica, Buffalo Grove, IL) with ×63 and ×100 magnification.

### Chromatin immunoprecipitation (ChIP) assay

ChIP assay was performed as described earlier (43,51) . Briefly, podocytes were treated with or without FG4592 and after incubation period, cells were cross linked by glutaraldehyde and these crosslinks are revealed by glycine. The fragment size was 500bp and incubated with protein A beads/ ZEB2 antibody and pull down was purified and subjected to RT-qPCR. Primers used in this experiment is: Forward primer 5′-GACCTAAGAGTCCACATCTG-3′ and Reverse primer 5′-CACAACTAGCCCAGGATAC-3′.

### Fluo3-AM Staining for Ca^2+^ influx assay

Intracellular Ca^2+^ levels were measured in podocytes by staining calcium with Fluo-3AM dye as previously described (52). To brief, human podocytes were cultured in 6-well plates at 1×10^6^ cells/well then treated with or without 2-APB for 2 h. After cells were treated with FG4592 for 4h, the Fluo3-AM dye was loaded into the wells at final concentration 4uM for 30 min in CO_2_ incubator at 37 °C. Cell lysates were prepared in calcium free HBSS solution and centrifuged at 400 × g for 5 min. Supernatant was assessed to measure fluorescence at λ485nm (excitation); λ538 (emission). The fluorescence maximum (Fmax) was achieved by adding 1% NP-40 to release maximum calcium bound dye from cells and EGTA (0.5 M) was added to cell lysate to quench calcium and considered this as fluorescence minimum (Fmin). Intracellular free calcium was quantified by the formula ; [Ca2+] = Fbasal – (Kd × ((F ™ Fmin)/(Fmax ™ F))) where Kd for Fluo3-AM is 390 nM. The data expressed as relative calcium fold change.

### Measuring TG2 activity

Trasglutaminases activity was performed in podocytes as previously described (53). Transglutaminase activity utilizes the deamidation reaction of the transglutaminase enzyme with a donor and acceptor substrate resulting in the formation of a hydroxamate product. In this assay, CBZ-Gln-Gly an amine donor and hydroxylamine as an amine acceptor were used for the transglutamination reaction. In the presence of calcium and glutathione, TG2 carries out deamidation reaction and forms CBZ-glutaminyl-glycyl-hydroxamate and can be measured at 525nm in spectrophotometer. Briefly, the assay was carried out by incubating 100ug of total cell lysates containing no CaCl_2_ with 0.23mL incubation solution containing 31 mM CBZ-L-glutaminyl-glycine, 174 mM Tris, 4 mM CaCl_2_, 8.7 mM glutathione (GSH) and 87 mM hydroxylamine at final concentrations. After a 10 min of incubation at 37°C, 0.5 mL of 12% V/V TCA was added to precipitate the protein and substrate complexes for 5 min. After high-speed centrifugation the final clear supernatant was measured at 525nm. The TG2 activity is expressed in units/mg of protein catalyze the formation of 1.0 µmole of hydroxamate per minute at pH 6.0 at 37°C.

### Transfection of Plasmid DNA and siRNA

HEK293T cells were cultured in DMEM containing 10% FBS and antibiotic & antimycotics. Before the day of transfection HEK293T cells were seeded as monolayers at 1×10^5^ cells/well in 6-well plates and grown overnight at 37°C in 5% CO_2_ incubator. Cells were subjected to transfection according to the manufacturer protocol with ZEB2 expression plasmid using X-tremeGENE 9 DNA Transfection Reagent and ZEB2 siRNA and TRPC6 siRNA using X-tremeGENE siRNA Transfection Reagent. The X-tremeGENE 9 DNA Transfection Reagent with DNA and X-tremeGENE siRNA Transfection Reagent with siRNA were mixed with serum free media incubated for 20min. After 20 min, this transfection complex was added to cells. After 6h, cells were treated with 10µM FG4592 for 24h. Cell lysates were analyzed by western blotting after transfection completion.

### Animal handling and treatment procedures

In this study we have used the 6-week old Swiss albino male mice weighing nearly 30±5g. The mice were randomly assigned to four groups (n=6): 1) control group, 2) FG4592 treated group, and 3) FG4592+Metformin treated group, 4) Metformin treated group. Experimental mice received a single i.p dose of FG4592 (5mg/kg/day) for everyday whereas control mice have received an equal volume of PBS for three months. The FG4592+Metformin treated group were received metformin (250mg/kg/day) per day through i.p, 3h prior to FG4592 treatment whereas metformin treated group received only metformin. After completing the experimental period, the mice urine were collected to examine the proteinuria and to estimate albumin and creatinine levels. After analyzing the urine, we have sacrificed the animals for further experiments. Institutional Animal Ethics Committee of the University of Hyderabad, approved the experimental procedures of animal study.

### Kidney samples from humans

Stored kidney biopsies without patient identification were obtained from Guntur General Hospital, Guntur, Andhra Pradesh, India. The study (GMC/IEC/120/2018) was approved by the Guntur Medical College and Government General Hospital’s Institutional Review Board. The principles of the Declaration of Helsinki are followed throughout the research.

### Histological Examination

Paraffin sections (4µM thickness) were used for H&E, PAS, and Masson trichrome staining. Images were acquired using a trinocular microscope (Leica, Buffalo Grove, IL). Renal cortex from control, FG4592 and FG4592 along with metformin treated tissue samples were fixed in 2.5% glutaraldehyde for 24 h, and TEM analysis performed as described earlier (42,43).

### Assessment of glomerular injury score

We performed PAS staining to quantify glomerular injury, such as mesangial matrix deposition, glomerular capillaries aberrations, and glomerular tuft area. We analyzed from each mouse 20 glomeruli sections in a double-blinded fashion by using ImageJ software, and the average value from each mouse was presented as a data point in the plot.

### Statistical analysis

The data are presented as mean ± S.E. of at least 3 independent experiments. Prism software (GraphPad Software Inc.) was used to analyze the data. Statistical differences between the groups made using Student’s t test and one way ANOVA. Statistical significance was determined as *p* < 0.05.

## DATA AVAILABILITY STATEMENT

The data that support the findings of this study are available on request from the corresponding author.

## ACKNOWLEDGMENTS

We acknowledge the funding from Indian Council of Medical Research (#2019-0905) and MoHFW-Department of Health Research (#12020/02/2019-HR) to AKP. Authors thank Dr. Rajkishor Nishad and Dr. Manga Motrapu for helpful discussion during the study and preparation of images. Authors thank Syed V. Tahaseen for her help with human kidney sections and urine analysis.

## AUTHOR CONTRIBUTIONS

AKP conceptualized the study. LPK, AKS, and RK performed experiments. LPK and AKP analyzed the data. LPK and AKS performed statistical analysis. LPK & AKP prepared the manuscript with inputs from AKS & RK.

## CONFLICT OF INTERESTS

The authors declare that there are no conflict of interests.

## Notes

### Competing Interest Statement

The authors have declared no competing interest.

